# The electron transport chain sensitises *Staphylococcus aureus* and *Enterococcus faecalis* to the oxidative burst

**DOI:** 10.1101/165589

**Authors:** Kimberley L. Painter, Alex Hall, Kam Pou Ha, Andrew M. Edwards

## Abstract

Small colony variants (SCVs) of *Staphylococcus aureus* typically lack a functional electron transport chain and cannot produce virulence factors such as leukocidins, hemolysins or the anti-oxidant staphyloxanthin. Despite this, SCVs are associated with persistent infections of the bloodstream, bones and prosthetic devices. The survival of SCVs in the host has been ascribed to intracellular residency, biofilm formation and resistance to antibiotics. However, the ability of SCVs to resist host defences is largely uncharacterised. To address this, we measured survival of wild-type and SCV *S. aureus* in whole human blood, which contains high numbers of neutrophils, the key defense against staphylococcal infection. Despite the loss of leukcocidin production and staphyloxanthin biosynthesis, SCVs defective for heme or menquinone biosynthesis were significantly more resistant to the oxidative burst than wild-type bacteria in a whole human blood model. Supplementation of the culture medium of the heme-auxotrophic SCV with heme, but not iron, restored growth, hemolysin and staphyloxanthin production, and sensitivity to the oxidative burst. Since *Enterococcus faecalis* is a natural heme auxotroph and cause of bloodstream infection, we explored whether restoration of the electron transport chain in this organism also affected survival in blood. Incubation of *E. faecalis* with heme increased growth and restored catalase activity, but resulted in decreased survival in human blood via increased sensitivity to the oxidative burst. Therefore, the lack of functional electron transport chains in SCV *S. aureus* and wild-type *E. faecalis* results in reduced growth rate but provides resistance to a key immune defence mechanism.

## Introduction

*Staphylococcus aureus* is responsible for a raft of different infections of humans and animals [1-3]. The key host defence against infection is the neutrophil, which phagocytoses *S. aureus* and exposes it to a cocktail of reactive oxygen species (ROS) during a process known as the oxidative (or respiratory) burst [4-6]. Whilst this is often sufficient to clear infection, invasive staphylococcal diseases frequently lead to persistent or recurrent infections of the bones, joints, heart or implanted devices [1, 7–9]. The development of these hard to treat infections is often associated with the presence of small colony variants (SCVs) [10–17]. As the name suggests, SCVs form small colonies on agar plates, typically due to metabolic defects caused by mutations that abrogate the electron-transport chain or biosynthetic pathways [16–21]. For example, several clinical studies have isolated SCVs with mutations in genes required for heme or menaquinone biosynthesis, including from the bloodstream [17–20]. The slow growth of SCVs provides a strong selection pressure for reversion to the wild-type, either by repair of the causative mutation or the acquisition of a suppressor mutation [18,19,22]. This presents challenges to their study and so targeted deletion of genes within the *hem* or *men* operons, which confer a phenotype that is identical to that of clinical SCVs, has been used to enable their study without the problem of reversion to the wild-type [23–26], SCVs can also arise in the absence of mutation, resulting in a very unstable phenotype, although the molecular basis for this is unknown [27], The emergence of SCVs is a rare but consistent consequence of *S. aureus* replication, which generates a small sub-population of the variants [22]. However, SCV emergence is significantly increased in response to diverse environmental stresses including antibiotics, reactive oxygen species, low pH within host cell vacuoles and exoproducts from *Pseudomonas*, which frequently causes co-infections with *S. aureus* [26–33].

Despite their diverse molecular basis, most SCVs have similar phenotypic characteristics. For example, activity of the Agr quorum-sensing system is weak or absent, and therefore cytolytic toxin production is negligible whilst surface proteins are strongly expressed [25,34–36]. These properties enable SCVs to persist in non-immune host cells and form robust biofilms, which has been hypothesised to contribute to their ability to persist in host tissues [27,37–39]. Furthermore, SCVs are typically resistant to antibiotics including the aminoglycosides, sulphonamides or fusidic acid and are often less susceptible to other antibiotics compared to wild-type bacteria [40–44].

Whilst these phenotypic properties very likely contribute to staphylococcal persistence in the host, the ability of SCVs to resist phagocytic cells, the key host defence against *S. aureus*, is poorly understood. Although SCVs are resistant to the ROS H_2_O_2_, they lack several defences used by wild-type bacteria to protect against immune cells [26]. For example, staphyloxanthin pigment, which promotes wild-type survival of both the oxidative burst and antimicrobial peptides, is absent in SCVs [15,18,45–47]. Furthermore, wild-type bacteria secrete numerous cytolytic toxins that kill neutrophils and enable bacterial survival, but this is absent in SCVs [15,18,25,34]. SCVs also exhibit reduced coagulase activity and some isolates lack catalase, both of which have been linked to survival of wild-type bacteria in the host [15,18,26,48–50]. Therefore, the effect of a defective electron transport chain on the susceptibility of SCV *S. aureus* to the oxidative burst of neutrophils is unclear.

*Enterococcus faecalis*, another major cause of bloodstream infections, shares some of the phenotypic properties of *S. aureus* SCVs since it is naturally defective for heme production and therefore lacks a functional electron-transport chain [51–53]. However, *E. faecalis* encodes type *a* and *b* cytochromes, and the presence of exogenous heme promotes *E. faecalis* growth in air, confirming the presence of an otherwise intact respiratory chain [51–53]. Exogenous heme also restores catalase activity, which has been shown to promote H_2_O_2_ resistance [54–55]. As such, it is unclear what whether *E. faecalis* gains an advantage from being defective for heme biosynthesis, particularly with respect to host defences that generate reactive oxygen species such as neutrophils.

Therefore, the aim of this work was to determine how the absence of the electron transport chain affects the survival of *S. aureus* and *E. faecalis* exposed to the oxidative burst of neutrophils.

## Methods

### Bacterial strains and culture conditions

The bacterial strains used in this study are detailed in table 1. Staphylococci were grown in tryptic soy broth (TSB) at 37 °C with shaking (180 RPM) for 18 h to late stationary phase. Enterococci were grown in Todd-Hewitt broth supplemented with 0.5% yeast extract (THY) at 37 °C with shaking (180 RPM) for 18 h to late stationary phase. For assays involving human blood, bacteria were plated onto Columbia blood agar (CBA) or THY supplemented with 5% sterile defibrinated sheep’s blood to neutralise any remaining oxidants from the assay. For some experiments iron (and other cations) was removed from TSB (100 ml) by incubation with Chelex resin (6 g) for 16 h at 4 °C with stirring. The following individual metals were then replaced: ZnCI_2_ (25 µM), CaCl_2_ (1 mM), MgCl_2_ (1 mM), MnCl_2_ (25 µM). Iron was added in the form of FeCl_3_ (1 or 10 µM) or heme (10 µM).

**Table 1.**
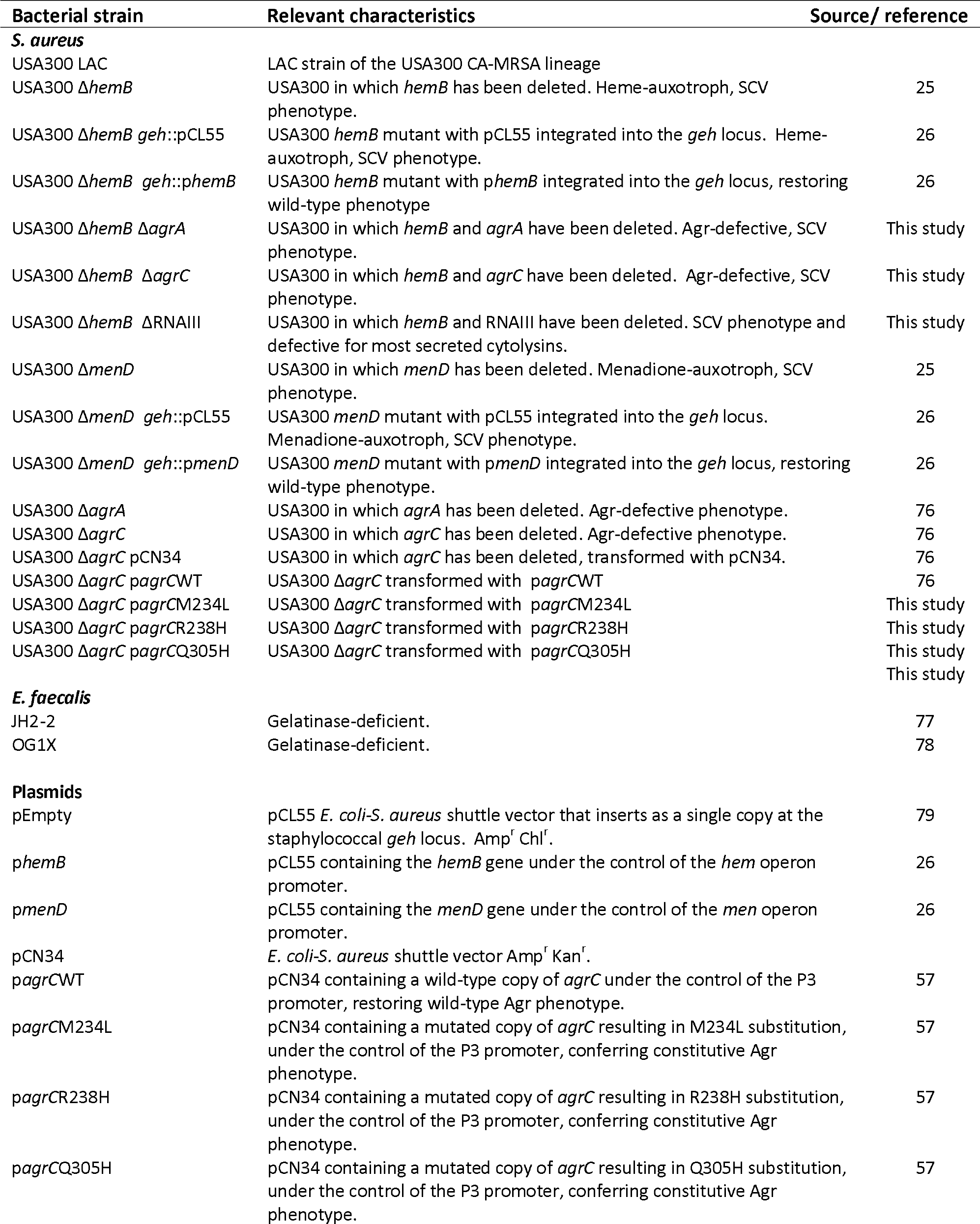
Bacterial strains and plasmids used in this study.

### Genetic manipulation of *S. aureus*

The construction of Δ*menD*, Δ*hemB* Δ*agrA*, Δ*agrC* and ΔRNAIII mutants was achieved using pIMAY as described previously [25,26,56]. To construct the double Δ*hemB*Δ*agr* mutants, the three *agr* mutants (Δ*agrA*, Δ*agrC* and ΔRNAIII) were made electrocompetent and the *hemB* gene deleted using pIMAY as described previously [25].

Mutants lacking Δ*hemB* or Δ*menD* were complemented with pCL55 containing the relevant gene under the control of the *hem* or *men* operon promoters respectively [26]. To control for pleiotropic effects of plasmid insertion into *geh*, pCL55 alone was transformed into *hemB* and *menD* mutant strains. The Δ*agrC* mutant was complemented with pCN34 containing a copy of the *agrC* gene under the control of the *agr* P3 promoter, and pCN34 alone (pEmpty) was used to control for pleitropic effects of the plasmid. In addition to wild-type *agrC*, plasmids containing mutated forms of *agrC* which confer a constitutively active phenotype were also transformed into the Δ*agrC* mutant strain [57].

### Whole human blood survival assay

The survival of bacteria in whole human blood was done as described previously [58]. Ethical approval for drawing and using human blood was obtained from the Regional Ethics committee and Imperial NHS trust tissue bank (REC Wales approval: 12/WA/0196, ICHTB HTA licence: 12275). Blood was drawn from healthy human donors into tubes containing EDTA and used immediately in assays based on a previously described protocol [4]. Suspensions of bacteria (10^5^ CFU in 10 µl PBS) were mixed with blood (90 µl) and incubated for up to 6 h at 37 °C with mixing. At indicated time points aliquots were taken, diluted serially in PBS and plated onto CBA plates to enumerate CFU. In some assays blood was pre-treated (10 min) with Diphenyleneiodonium (DPI) or an identical volume of DMSO alone to control for solvent effects [4].

### Measurement of bacterial growth

Stationary-phase bacteria were diluted 1:50 into a final volume of 200 µl TSB in microtitre plates (Corning) before incubation at 37 °C with shaking (500 RPM) in a POLARstar Omega multiwell plate reader. Bacterial growth was measured using OD_600_ measurements every 30 min for a total of 17 h [57].

### Hemolysin production

The hemolytic activity of bacterial culture supernatants was determined as described previously [25]. Briefly, culture supernatants were recovered by centrifugation (13,000 X *g*, 10 min) of stationary-phase cultures. The supernatant was then diluted in 2-fold steps using fresh TSB. Aliquots from each dilution (100 µl) were mixed with an equal volume of 2 % sheep blood suspension in PBS and incubated at 37 °C for 1 h in a static incubator. Subsequently, unlysed blood cells were removed by centrifugation and the supernatant containing lysed erythrocytes transferred to a new microtitre plate. The degree of erythrocyte lysis was quantified by measuring the absorbance of the supernatant at A_450_ and reference to controls. Erythrocytes incubated with TSB alone or TSB containing 1 % TX-100 served as negative and positive controls respectively.

### Measurement of phagocytosis and immune cell viability

Phagocytosis of bacteria in whole human blood was determined using a protocol based on that described previously [59]. Stationary-phase bacteria (1 ml) were pelleted (17,000 X *g*, 3 min) and washed twice with PBS. The pellet was then resuspended in 200 µl of 1.5 mM Fluorescein isothiocyanate (FITC) dissolved in freshly prepared carbonate buffer (0.05 M NaCO_3_ and 0.1 M NaCl). Bacteria were then incubated for 60 min (room-temperature with tumbling) in the dark. FITC-labelled bacteria were then washed three times in carbonate buffer and adjusted to 1 × 10^6^ CFU ml^-1^ in PBS. FITC-labelled bacteria (10 µl, 1 × 10^4^ CFU) were added to 96-well plates prior to the addition of 90 µl of freshly isolated blood, as described for the whole blood killing assay. At each time point (0, 2, 4 and 6 h), the blood/bacteria mixture (100 µl) was added to 900 µl red blood cell lysis solution (eBioscience) and incubated at room temperature in the dark for 10 min. Samples were then centrifuged (500 × *g*, 10 min) and the resulting pellet washed once in PBS (1 ml) before a final centrifugation step (500 × *g*, 10 min) and then the pellet containing immune cells and bacteria was resuspended in 100 µl PBS or 1% paraformaldehyde (PFA; Affymetrix) if no further staining was required. Where samples were to be analysed for host cell death, samples were incubated in PBS containing the Zombie Violet live-dead dye (Biolegend) at a 1:500 dilution in the dark. Free primary amine groups were quenched using 1.4 ml 1% bovine serum albumin (BSA) and samples were centrifuged (500 × *g*, 10 min) before resuspension in 100 µl 1% PFA. Positive controls were generated by heat-killing host cells (100 °C, 10 min) prior to Zombie staining. Samples were then fixed overnight (12-16 h) in 1% paraformaldehyde at 4 °C. Immune cell/bacteria samples were analysed on a Fortessa flow cytometer (BD) and at least 10,000 events were captured. Green (FITC-bacteria) and violet (Zombie-labelled host cells) fluorescence were detected at 488/530 (30) nm and 404/450 nm, respectively. Based on preliminary analyses and using the methodology of Surewaard *et al*. (2013) [60], free bacteria (i.e. bacteria not phagocytosed) were identified as events with a side scatter of < 50K. By contrast, host cells were identified as events with a side scatter of > 50K. Samples were analysed alongside controls, which consisted of bacteria without FITC labelling, host cells with or without Zombie stain, uninfected host cells and heat-killed host cells as appropriate. Data were analysed using FlowJo software (Version 10). Compensation was not necessary as the spectra of the fluorescent signals did not overlap.

### Catalase assay

Catalase activity of bacterial cells was determined as described previously [26]. Overnight bacterial cultures (1 ml) were washed three times in PBS and 10^7^ CFU added to 100 µM H_2_O_2_ in PBS (1 ml). Bacteria were incubated in the H_2_O_2_ in the dark at 37 °C. At the start of the assay and every 15 min, 200 µl of sample was pelleted (17,000 × g) and 20 µl added to a 96 well microtitre plate. The concentration of remaining H_2_O_2_ was determined using a Pierce Quantitative Peroxide Assay (Aqueous Compatible) kit.

## Results

### The loss of the electron transport chain promotes survival of *S. aureus* in human blood

To study the susceptibility of electron transport chain-deficient SCVs to the oxidative burst, we employed the well-established *ex vivo* whole human blood model of infection. This is model is appropriate because *S. aureus* is a major cause of bacteraemia and blood contains a high density of neutrophils, as well as the required opsonins and other relevant immune factors such as platelets [4,61,62]. In this model system, *S. aureus* is rapidly phagocytosed by neutrophils and exposed to the oxidative burst [4,61,62].

Freshly-drawn human blood containing anti-coagulant (EDTA) was incubated with wild-type *S. aureus* USA300, or mutants with deletions of *hemB* or *menD*, and survival determined over time by CFU counts. Preliminary experiments determined that individual donors had slightly different anti-staphylococcal activity and so at least 3 different donors were used for each experiment (Fig. 1A). However, for each of the 5 donors we observed a consistent decrease in CFU counts of wild-type bacteria over time with just 1-5% of the inoculum surviving after 6 h (Fig. 1A). By contrast, SCVs defective for heme-or menaquinone-biosynthesis survived at much higher levels than the wild type over the entire duration of the assay with 70% of the Δ*hemB* mutant inoculum and 69% of Δ*menD* viable after 6 h incubation in blood (Fig. 1B,C).

**Figure 1.**
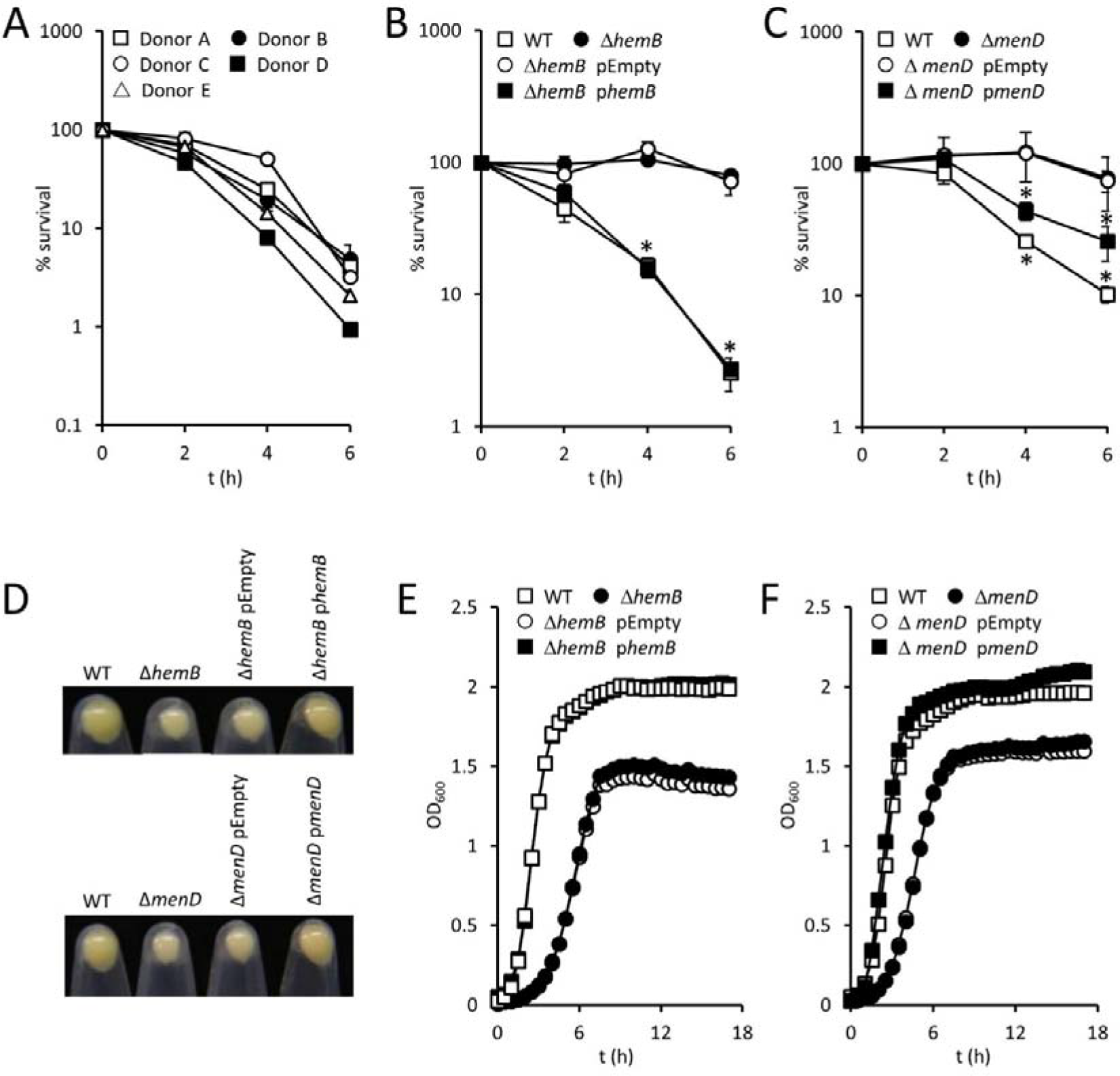
Survival of SCV *S. aureus* in blood is greater than that of wild-type bacteria. (A) Survival of wild-type *S. aureus* USA300 in blood from individual donors. Data represent the mean survival from 3 independent experiments from each donor. (B, C) Survival of wild-type *S. aureus* USA300 and Δ*hemB* (B) or Δ*menD* (C), and complemented strains, in human blood. Data represent the mean of 4 independent experiments using blood from at least 3 different donors. (D) Images of pelleted stationary-phase *S. aureus* strains highlighting differences in pigmentation. Images are representative of 3 independent assays. (E, F) Growth profiles of *S. aureus* wild-type and Δ*hemB* (E) or Δ*menD* (F), and complemented strains. Data represent the mean of 3 independent experiments. Where shown, error bars represent the standard deviation of the mean. Data in B and C were analysed by 2-way repeated measures ANOVA and Dunnett’s post-hoc test. *represents p = <0.01 compared with wild-type.

Complementation of the *hemB* or *menD* mutations conferring the SCV phenotype restored the wild-type phenotype for growth and staphyloxanthin production, and resulted in significantly decreased survival in blood (Fig. 1B,C,D,E,F). This confirmed that enhanced SCV survival in blood was due to the loss of heme or menaquinone biosynthesis, rather than the acquisition of adventitious mutations during genetic manipulation. Therefore, despite the lack of staphyloxanthin pigment and cytolysin production, loss of the electron transport chain confers a survival advantage to *S. aureus* in blood.

### Wild-type *S. aureus* is more sensitive to the oxidative burst than SCVs

Having demonstrated that survival of SCVs in blood is greater than that of the wild-type, we sought to understand why. Several previous studies have shown that incubation of *S. aureus* in whole human blood results in rapid phagocytic uptake of the bacterium by polymorphonuclear leukocytes (PMNs) [4,61,62]. We confirmed those findings and found no differences in the phagocytosis of wild-type, Δ*hemB* or Δ*menD* mutants (Fig. 2A). We also demonstrated that the viability of neutrophils that phagocytosed *S. aureus* did not vary between wild-type and SCVs (Fig. 2B). Therefore, both immune evasion and killing of immune cells by SCVs were ruled out as an explanation for their ability to survive in human blood.

**Figure 2.**
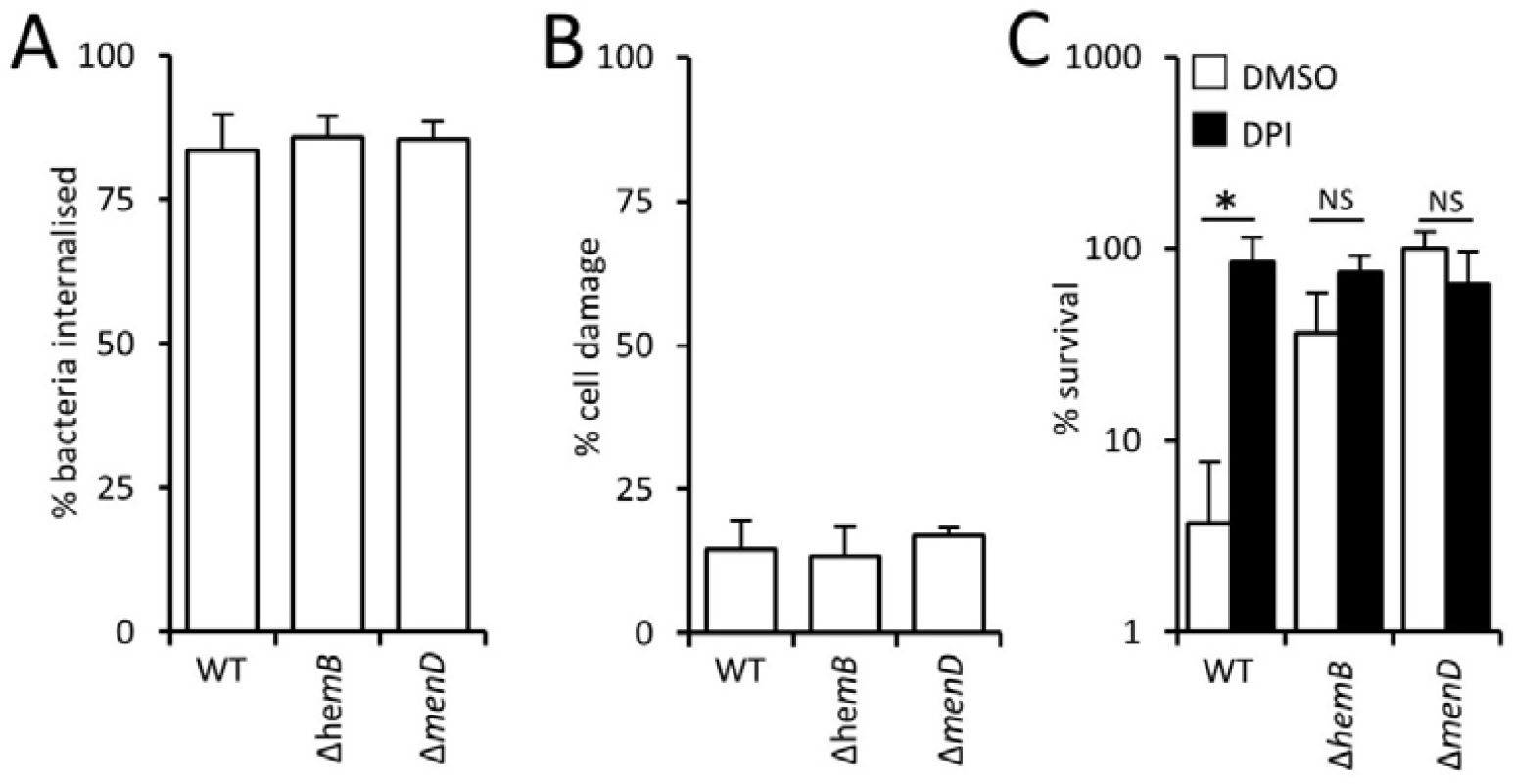
SCVs survive the oxidative burst better than wild-type *S. aureus*. (A) The percentage of *S. aureus* wild-type, Δ*herriB*, or Δ*menD* 5. *aureus* USA300 internalised into phagocytic cells 2 h after inoculation into whole human blood. (B) The percentage of phagocytic cells that contained *S. aureus* strains, and had impaired membrane integrity, as determined using the Zombie Violet reagent after 6 h in whole human blood. (C) Survival of *S. aureus* wild-type, Δ*hemB*, or Δ*menD S. aureus* USA300 after 6 h in blood pre-treated with the NADPH oxidase inhibitor diphenyleneiodonium (DPI) or an identical volume of DMSO solvent alone (DMSO). Data were analysed via a one-way ANOVA with Tukey’s post hoc test, which revealed no significant differences in (A) or (B). In (C), * indicates p = < 0.01, NS indicates p = > 0.05 when the indicated comparisons were made.

The principle mechanism by which neutrophils kill *S. aureus* is the oxidative burst [4–6]. To confirm that this was the case in our model system we measured bacterial viability in human blood treated with diphenyleneiodonium (DPI), which blocks the oxidative burst, or the DMSO solvent alone. Suppression of NADPH with DPI, but not DMSO alone, resulted in significantly elevated survival of wild-type S. *aureus*, confirming that the oxidative burst is the key defence against *S. aureus* in human blood (Fig. 2C) [4–6]. The addition of DPI to blood did not significantly alter SCV CFU counts, since survival was already very high (Fig. 2C). Therefore, SCV *S. aureus* appears to be significantly less susceptible to the oxidative burst than wild-type bacteria. This is in agreement with our previously reported finding that both the Δ*hemB* and Δ*menD* SCVs were more resistant to H_2_O_2_ than wild-type bacteria, and provides an explanation for the increased survival of SCVs in blood [26].

### Agr activity promotes the survival of wild-type but not SCV *S. aureus* in blood

Although Agr-regulated toxins have been shown to kill neutrophils, several clinical studies have shown an association of Agr dysfunction with persistent bacteremia [63]. Therefore, we considered the possibility that the weak Agr activity of SCVs contributed to their survival in blood.

To test this, we compared the survival of wild-type and Agr-defective strains in whole human blood. This revealed a significantly greater loss of viability of *agr* mutants compared with the wild-type (Fig. 3A). In particular, mutants lacking quorum-sensing components of Agr (Δ*agrA* or Δ*agrC)* were approximately 4-fold more susceptible to immune cells in blood than the wild-type, whilst the RNAIII mutant was 2-fold more susceptible than the wild-type (Fig. 3A). This finding is in keeping with previous work that showed that AgrA-regulated PSMs contribute to survival of *S. aureus* within the phagocytic vacuole of neutrophils, in addition to RNAIII-regulated toxins [60]. Therefore, a functional Agr system promotes the survival of wild-type bacteria in human blood.

**Figure 3.**
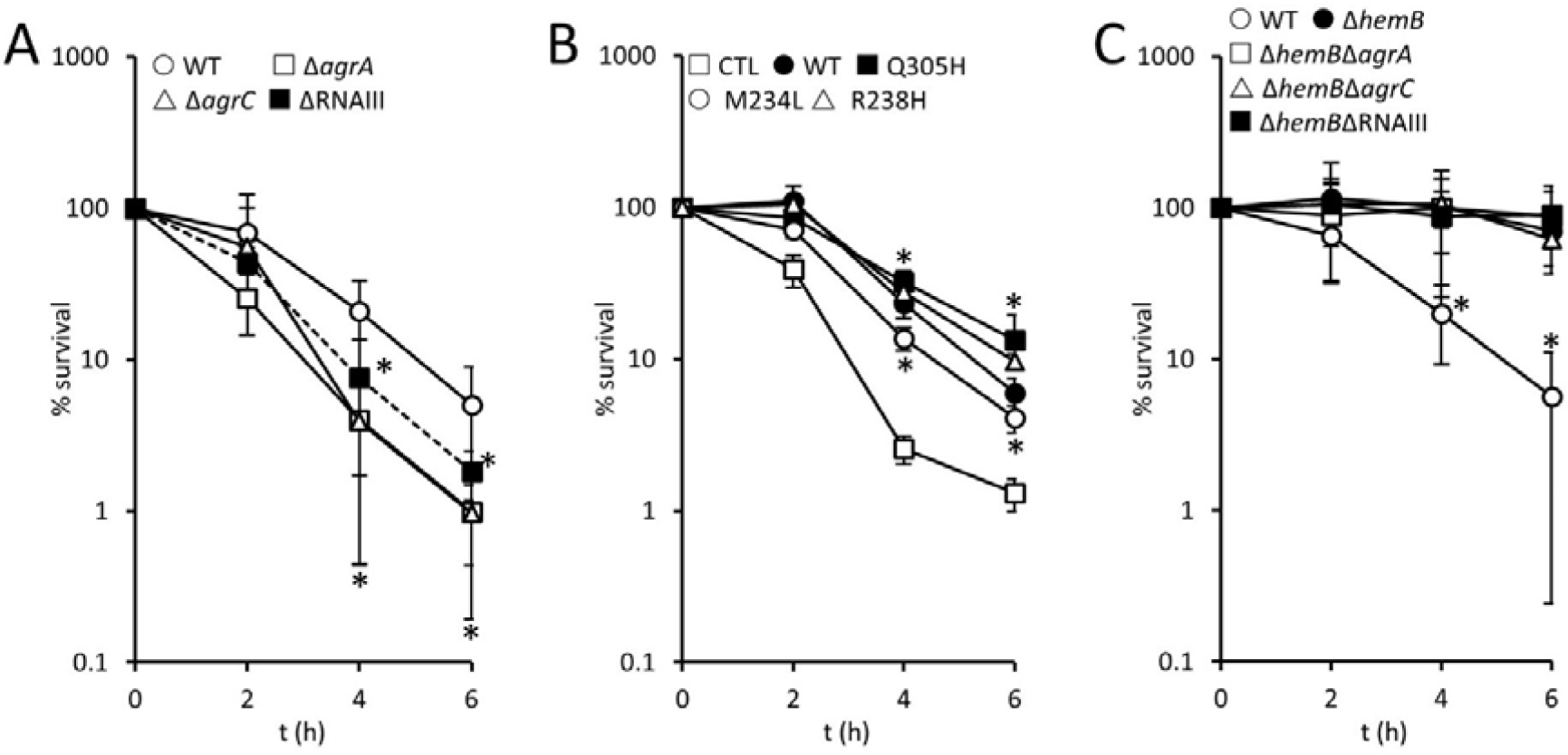
Survival of wild-type but not SCV *S. aureus* is enhanced by Agr. (A) Survival of wild-type (WT), Δ*agrA*, Δ*agrC* or ΔRNAIII *S. aureus* USA300 in whole human blood over 6 h. (B) Survival of *S. aureus* USA300 Δ*agrC* mutant transformed with pCL55 (CTL), pCL55 containing the wild-type *agrC* gene (WT), or 3 mutated variants of *agrC* that result in Q305H, M234L or R238H substitutions conferring a constitutively-active phenotype. (C) Survival of wild-type (WT), Δ*hemB*, Δ*hemB*Δ*agrA*, Δ*hemB*Δ*agrC* or Δ*hemB*ΔRNAIII *S. aureus* USA300 in whole human blood over 6 h. For all panels, data represent the mean of 4 independent experiments using blood from at least 3 different donors. Data were analysed by 2-way repeated measures ANOVA with Dunnett’s post-hoc test to compare strains to WT (A), CTL (B) or to Δ*hemB* (C). * indicates p = < 0.01. In panel A, all mutants were significantly more susceptible to immune defences than the wild-type at 4 and 6 h. In panel B, all strains expressing *agrC* (wild-type or mutated) survived better than the Δ*agrC* mutant at the 4 and 6 h time points. In panel C, all Δ*hemB* mutants (+/-*agr)* survived equally well and significantly better than the wild-type.

Complementation of the Δ*agrC* mutant with a wild-type copy of the gene increased survival in blood (Fig. 3B). However, complementation of Δ*agrC* with mutant copies of *agrC* which confer constitutive Agr activity, even in the presence of serum [57], did not promote bacterial survival above that of the wild-type gene (Fig. 3B).

Although Agr activity is extremely weak in SCVs, we explored whether this contributed to their survival by generating Δ*hemB* mutants defective for *agrA, agrC* or RNAIII, and measuring their survival in blood (Fig. 3C). This revealed that survival of each of the Δ*hemB*Δ*agr-mutants* was as high as for the Δ*hemB* mutant with an intact *agr* operon. Therefore, whilst loss of Agr activity in the wild-type reduces survival in human blood, the lack of Agr activity in SCVs is not detrimental for their survival. This indicates that toxin production is an important mechanism by which wild-type *S. aureus* survives phagocytosis. By contrast, since SCVs can survive the oxidative burst they do not need toxins to survive phagocytosis.

### Restoration of the electron transport chain with heme results in decreased survival of SCVs in blood

During infection, *S. aureus* acquires iron from the host, predominantly via the acquisition of heme liberated from erythrocytes via hemolytic toxins [64]. In addition to acting as an iron source, heme can also be utilised by heme-auxotrophic SCVs to restore the electron transport chain [18,26]. To determine how heme influenced the phenotype of heme- and menquinone-defective SCVs, and their susceptibility to the oxidative burst, we grew wild-type or SCV *S. aureus* in media deficient for heme and containing minimal free iron (1 µM FeCl_3_), abundant iron (10 µM FeCl_3_), or in the presence of heme (10 µM).

The growth rate of wild-type *S. aureus* was not significantly affected by the presence of the higher concentration of FeCI_3_ or heme, although the latter led to a slight increase in the length of the lag phase (Fig. 4A). Similarly, abundant iron did not affect growth of the Δ*menD* SCV, but heme caused slight growth retardation (Fig. 4A). By contrast, abundant iron slightly promoted the growth rate of the Δ*hemB* SCV, whilst heme enhanced the growth almost to wild-type levels (Fig. 4A). In addition to the growth rate, heme supplementation restored hemolytic activity and staphyloxanthin biosynthesis to the Δ*hemB* mutant (Fig. 4B). However, heme supplementation of the Δ*hemB* mutant also resulted in significantly increased susceptibility to the oxidative burst of neutrophils in blood (Fig. 4C), which is in keeping with our previous finding that heme supplementation renders heme-auxotrophic SCVs sensitive to H_2_O_2_ [26]. By contrast, supplementation of the medium with iron had no effect on susceptibility of the Δ*hemB* mutant to the oxidative burst or H_2_O_2_ (Fig. 4C). This is in agreement with previous work showing that iron-loading of *S. aureus* does not alter susceptibility to the oxidative burst of neutrophils [65,66]

**Figure 4.**
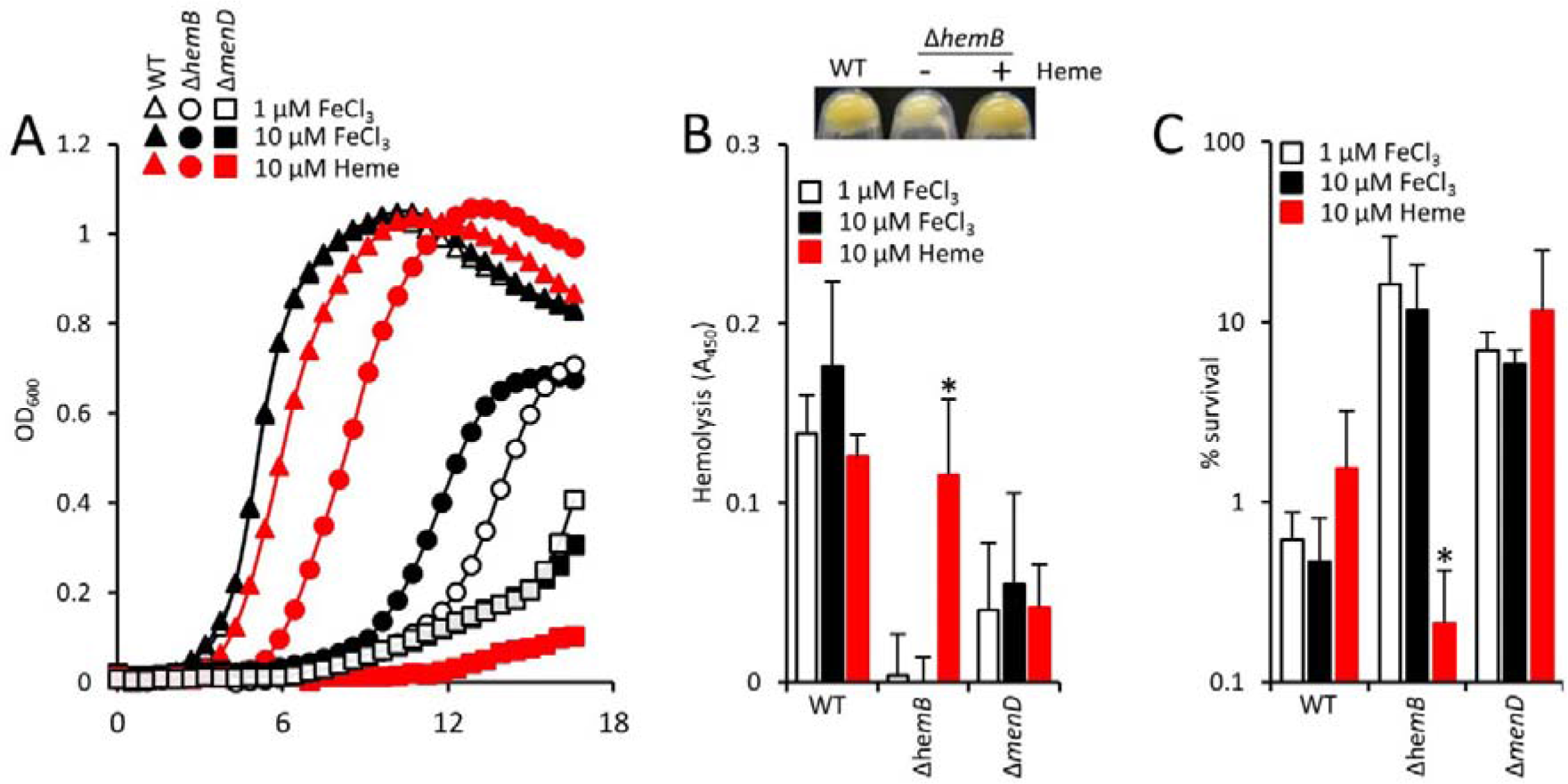
Heme promotes growth and virulence factor production of the Δ*hemB* mutant but decreases survival in blood. (A) growth profiles (as determined by OD_600_ readings) of WT, Δ*herriB* and Δ*menD* in metal-adjusted TSB containing iron in the form of 1 or 10 µM FeCl_3_ or 10 µM heme. Please note that the open triangles are largely obscured by the filled triangles. (B) the graph shows hemolytic activity of WT, Δ*hemB* and Δ*menD* grown in the presence of 1 or 10 µM FeCl_3_ or 10 µM heme. The panel illustrates the pigmentation of the Δ*hemB* mutant grown in the absence or presence of 10 µM heme. WT is shown for comparison. There was no effect of heme on the pigmentation of the WT or Δ*menD* strain. (C) Survival of WT, *&hemB* and *hmenD*, grown in the presence of 1 or 10 µM FeCl_3_ or 10 µM heme, after 6 h incubation in whole human blood. Data (B) and (C) were analysed via a one-way ANOVA with Tukey’s post hoc test. For each strain, comparisons were made between 1 µM FeCl_3_ and 10 µM FeCl_3_ or 10 µM heme. * indicates p = **<**0.01.

By contrast to the Δ*hemB* mutant, the susceptibility of both the wild-type and Δ*menD* mutant to the oxidative burst was unchanged by growth in the presence of heme. Therefore, at the concentration used (10 µM), heme does not directly sensitise *S. aureus* to the oxidative burst. Rather, it is the restoration of the electron-transport chain in the Δ*hemB* mutant that confers sensitivity to the oxidative burst.

### The absence of an electron-transport chain enables survival of *Enterococcus faecalis* in human blood

The elevated survival of the *S. aureus* Δ*hemB* mutant, relative to wild-type, led us to consider whether a similar phenomenon occurred with *Enterococcus faecalis*, which despite producing cytochromes lacks a functional electron transport chain due to an inability to synthesise heme [51–53]. However, *E. faecalis* employs heme uptake systems to scavenge heme from the environment and therefore supplementation of the culture medium with heme results in increased growth under aerobic conditions. We confirmed this in two different *E. faecalis* strains (Fig. 5A,B), which grew to a higher optical density in the presence of heme. In addition, *E. faecalis* grown in the presence of heme produce a functional catalase, which we observed in both of the strains examined (Fig. 5C,D). However, as observed for the Δ*hemB* SCV, growth of *E. faecalis* in the presence of heme led to significantly diminished survival in human blood by increasing sensitivity to the oxidative burst (Fig. 5E,F). Therefore, as for SCV *S. aureus*, the absence of the electron-transport chain in *E. faecalis* promotes survival in the bloodstream by reducing sensitivity to oxidative stress generated by host immune cells.

**Figure 5.**
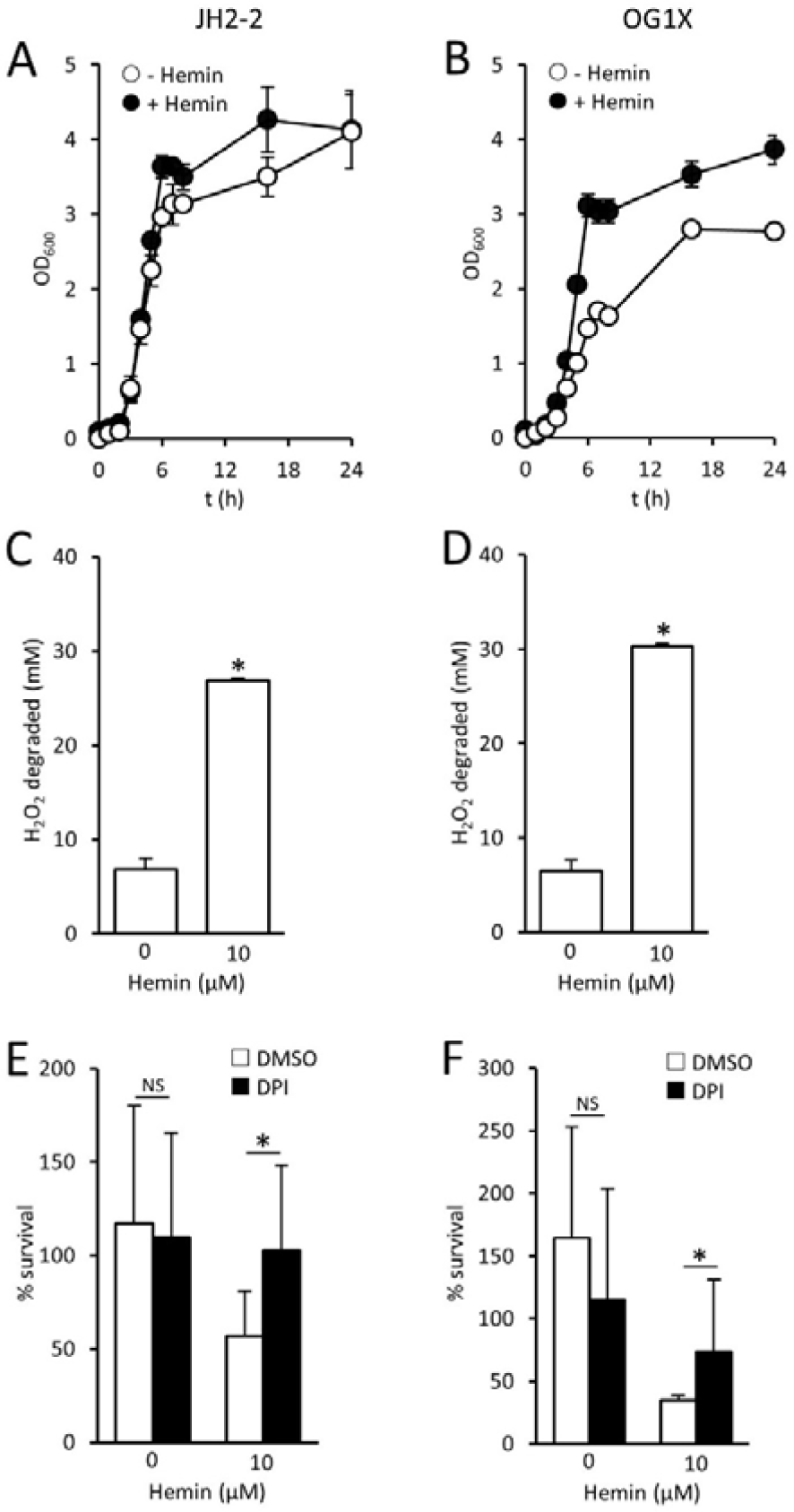
Heme promotes susceptibility of *E. faecalis* to host defences. (A,B) growth profiles (as determined by OD_600_ readings) of *E. faecalis* JH2-2 (A) or OG1X (B) grown in the absence or presence of 10 µM heme. (C,D) Catalase activity (expressed as mM H_2_O_2_ degraded in 1 hr by 10^7^ CFU) of *E. faecalis* JH2-2 (C) or OG1X (D) grown in the absence or presence of 10 µM heme. (E,F) Survival of *E. faecalis* JH2-2 (E) or OG1X (F) after 6 h in blood pre-treated with DPI or an identical volume of DMSO solvent alone. Data were analysed by one-way ANOVA with Tukey’s post hoc test, which revealed no significant differences (p = < 0.01) in (A) and (B) between bacteria grown in the absence or presence of heme. In (E) and (F), * indicates p = < 0.01, NS indicates p = > 0.05 when the indicated comparisons were made.

## Discussion

During infection *S. aureus* faces two major threats: host defences and antibiotic therapy. Previous work has shown that SCVs of *S. aureus* are less susceptible to antibiotics than wild-type bacteria. Our data demonstrate that the SCV *S. aureus* is also less susceptible to host immune defences. These data fit with a previous study that revealed that SCVs are less sensitive than wild-type to host-derived antimicrobial peptides [67]. However, the resistance of SCVs to both the oxidative burst and AMPs is surprising given the lack of staphyloxanthin pigment, which contributes to resistance of wild-type *S. aureus* to both ROS and AMPs [4,68].

We do not currently understand the molecular basis of ROS resistance in SCVs. However, the damaging effects of ROS are proposed to occur via the Fenton reaction, which involves the reaction of H_2_O_2_ with free iron leading to the generation of highly-reactive hydroxyl radicals [69,70]. The lack of an electron-transport chain, together with the associated decreased TCA activity (which utilises iron-containing enzymes such as aconitase) in SCVs is hypothesised to result in decreased iron content relative to wild-type bacteria. Furthermore, there is evidence that the electron transport chain generates superoxide radicals that liberate iron from iron-sulphur clusters, making it available for the Fenton reaction [71].

The ability of *S. aureus* SCVs to survive the oxidative burst comes at a cost. The electron-transport chain enables aerobic respiration, rapid bacterial growth and toxin production. These toxins include hemolysins that enable *S. aureus* to access heme, the bacterium’s primary source of iron during infection [72]. Therefore, the absence of hemolysin production by the Δ*hemB* mutant enables maintenance of the SCV phenotype in the presence of red blood cells. The menaquinone-defective SCV cannot restore the wild-type phenotype using host-derived materials and therefore maintains its phenotype regardless of hemolysin production.

*E. faecalis* lacks the necessary biosynthetic machinery to synthesise heme making it a heme auxotroph [51–53]. However, some strains secrete a cytolysin with hemolytic activity that provides a mechanism of heme acquisition[73]. The liberation of heme from erythrocytes would be expected to promote growth and restore catalase activity, but would also increase susceptibility to host defences. The maintenance of cytochromes and catalase that are restored by exogenous heme suggests that heme acquisition is a consistent and beneficial event during colonisation and/or infection. What is not clear however, is when and where heme acquisition occurs. For example, isolates recovered from patients with infective endocarditis, an infection of the heart valves that persists despite a robust immune response, are typically defective for hemolysin production [73,74]. This may indicate that hemolysin production, and thus heme acquisition, is undesirable at this site. By contrast, 30-40% of *E. faecalis* isolates carried in the gut or isolated from urinary-tract infections are hemolytic [74]. However, further work is needed to understand the basis for this observation and whether heme-mediated susceptibility to the oxidative burst plays a role.

Previous work reported that heme supplementation enabled *E. faecalis* to survive H_2_O_2_ challenge by restoring catalase activity [54–55]. However, whilst we also observed restoration of catalase activity in *E. faecalis* supplied with heme, this did not correlate with increased resistance to the oxidative burst. We have shown previously that the Δ*hemB* mutant is catalase-deficient but was much less susceptible to the oxidative burst than wild-type bacteria [26]. This indicates that catalase is not required for survival of the oxidative burst, a finding that is supported by previous work with *S. aureus* that showed a catalase mutant was as virulent as the wild-type [75].

In summary, SCV *S. aureus* sacrifices fast growth and toxin production for enhanced resistance to host defences and antibiotics. This dramatic change in phenotype may enable the transition from highly-damaging, acute infection to a less pathogenic but persistent infection type. Our data indicate that the lack of heme production in *E. faecalis* also promotes survival in human blood, suggesting a common survival mechanism between these two pathogens.

## Acknowledgements

The following are gratefully acknowledged for providing bacterial strains, phage or reagents: Ruth Massey (University of Bath), Malcolm Horsburgh (University of Liverpool), Tim Foster (Trinity College Dublin), Angela Nobbs (University of Bristol), and the Network on Antimicrobial Resistance in *Staphylococcus aureus* (NARSA) Program: under NIAID/NIH Contract No. HHSN272200700055C. A.M.E. acknowledges funding from the Royal Society, Department of Medicine (Imperial College), and from the Imperial NIHR Biomedical Research Centre, Imperial College London. K.L.P. was supported by a PhD studentship from the Faculty of Medicine, Imperial College London. K.P.H. is supported by an MRC-funded PhD studentship awarded to the Centre for Molecular Bacteriology and Infection, Imperial College London.

